# Injury caused by alcoholic cardiomyopathy in spontaneous ethanol drinking rats

**DOI:** 10.1101/2021.03.12.435159

**Authors:** Victória Mokarzel de Barros Camargo, Vanessa Caroline Fioravante, Patricia Fernanda Felipe Pinheiro, Francisco Eduardo Martinez

**Affiliations:** Department of Structural and Functional Biology, Institute of Biosciences of Botucatu, UNESP - Univ Estadual Paulista, Botucatu, SP, Brazil

**Keywords:** alcoholic cardiomyopathy, heart, ethanol, UChB rats, alcoholism

## Abstract

When speaking of pathologies caused or aggravated by the constant ingestion of ethanol, people with liver and central nervous system diseases soon come to mind, however, the acute intake of large amounts of ethanol and chronic abuse induce toxic effects in the majority of tissues. The heart is highlighted, since alcoholic cardiomyopathy (AC) has prevalence among alcoholics of 23 to 40% and occurs more frequently in men than in women. AC is characterized by dilation and poor contraction of one or both ventricles in the presence of increased ventricular wall thickness, along with a long history of ethanol abuse and no other cause identified. Our aim is to quantify the rate of cardiac tissue replacement, collagen fiber deposition and pro inflammatory cytokines in the left ventricle myocardium of volunteer ethanol drinking rats.

## 1. INTRODUCTION

Ethanol has been present in the daily life of humans since prehistory, when the first alcoholic beverages appeared as well as the appearance of agriculture and the invention of ceramics [1]. Ethanol is responsible for the death of approximately 2.5 million people per year worldwide, according to the World Health Organization [2]. It occupies third place in the mortality rate, with more than 50% of these deaths being caused by external causes and almost a third of cardiovascular deaths occur in the context of excessive drinking [3]. When talking about pathologies caused or aggravated by the constant ingestion of ethanol, illnesses linked to the liver and the central nervous system (CNS) come to mind, however, the acute ingestion of large amounts of ethanol and chronic abuse induce effects toxic in most tissues [4].

Among these organs, the heart stands out, since alcoholic cardiomyopathy (AC) has a prevalence, among alcoholics, of 23 to 40% and occurs more frequently in men than in women [5]. This disease is characterized by deficient dilation and contraction of one or both ventricles in the presence of increased ventricular wall thickness, along with a long history of ethanol abuse with no other identified cause. In histological evaluation, dilation, myofibrillar necrosis and fibrosis are typically present, with a reduction in myofibrils and giant mitochondria [4]. One of the possible explanations involves the inflammation of cardiomyocytes caused by ethanol and its metabolites. There is a link between chronic ethanol consumption and the production of pro-inflammatory cytokines, subsequent activation of inflammatory signaling pathways in the heart and CNS [6]. Consequently, there is oxidative stress and degenerative processes.

There is a loss of cardiomyocytes due to apoptosis owing to high concentrations of ethanol in animals. Also, cases of AC present cardiomyocytes with moderate hypertrophy, cell death, and dilated muscle [7]. Our experimental model is the variety of Wistar UChB rats which ingest ethanol spontaneously and chronically. The lineage emerged at the University of Chile through the inbreeding of rats that voluntarily ingested high amounts of ethanol [8]. The model is similar to the alcoholic man in several aspects, which allows data obtained in animals to direct clinical findings.

Some mechanisms try to explain the decrease in the number of cardiomyocytes in AC: imbalance in the rate of cardiac tissue replacement (relationship between mitosis and cell death), direct tissue damage (with consequent deposition of dense connective tissue and healing process) and triggering of the chronic inflammatory process (increased pro-inflammatory cytokines). Thus, our goal is to quantify the rate of cardiac tissue replacement, the deposition of collagen fiber and the proinflammatory cytokines in the left ventricular myocardium of ethanol voluntary drinking rats.

## 2. METHODOLOGY

### 2.1 Animals

Ten adult male rats 115 days old (adults), weighing between 300-400 g, *Rattus norvegicus albinus*, of the Wistar variety, called UChB (University of Chile), spontaneous drinkers with a high concentration of ethanol (consumption greater than 2 g of ethanol / Kg body weight/day), and ten adult male UChB rats 130 days old, weighing between 300 - 400 g, not exposed to ethanol, called UChBC rats obtained from the Department of Anatomy, Institute of Biosciences, Botucatu Campus, UNESP - Paulista State University. The rats were individually housed in polypropylene cages (43X30X15 cm) with shavings and kept at constant room temperature (23 ± 1 ° C) and lighting (12 h light / dark cycle, with the lights on 6 h). The rats were treated with a solid diet consisting of standard rodent food and filtered water at will.

The experimental protocol (1185/19-CEUA) followed the ethical principles in animal research governed by Federal Law 11,794 of October 8, 2008, regulated by Decree No. 6,899 of 2009, of according to the resolutions the National Council for the Control of Animal Experimentation (CEUA).

### 2.2 Experimental design

When the rats reached 65 days of age, two bottles were given, one containing ethanol solution and the other water, free choice during 15 days. Rats with ethanol consumption> 2 g ethanol / kg / day were eligible for the study. Subsequently, the rats with the profile of the UChB variety were divided into two groups (n = 20): the UChB group, in which the rats had free access to a 10% (v / v) ethanol solution and water at ease for 50 days and the UChBC group, in which the rats did not have access to an ethanol solution, only water at ease for 50 days.

### 2.3 Collect

The animals were first placed in carbon dioxide saturation chambers to be anesthetized for the euthanasia procedure. Subsequently, the rats were decapitated to collect blood and the chest cavity was opened to collect the heart. The heart was sectioned in the median sagittal plane with the aid of a stainless steel blade (Feather brand) and its parts fixed in 4% paraformaldehyde for 24 hours. Subsequently, they were washed in running water for 72 h and stored in 70 % ethanol.

### 2.4 Histological analysis

We evaluated the integrity of the cardiac tissue structure stained with Hematoxylin & Eosin (HE) and the healing process, if present, by marking the bundles of collagen fibers with Masson’s Trichrome (MT). The fragments of the left ventricle of the heart were embedded in paraffin, sectioned to four μm thick and stained with HE and MT. The sections were examined under a light microscope.

### 2.5 Immunohistochemistry

To assess the imbalance in the rate of cardiac tissue replacement, tests related to mitosis (Ki67) and cell death due to apoptosis (Caspase 3) were performed. The tissue sections of the 10 rats in each group were stained with antibodies against Ki-67 (Novocastra, Laboratories Ltd) and Caspase 3 (Biocare - USA), according to the manufacturing protocol. After immune reactions, the slides were washed in a TBS-T buffer and incubated with secondary antibodies (Polymer Anti-Mouse IgG or Anti-Rabbit - DAKO & CYT) for 1 h. Then, the slides were stained with diaminobenzidine (DAB; Sigma, St. Louis, MO, USA) and contrasted with Hematoxylin. Five slides from each left ventricle with three sections per slide were stained. The sections were analyzed and photomicrographed using the Zeiss microscope, model Axiophot2 Plus, with a camera (AxioCam MR1) coupled to the microscope and analyzed by the program Axio Vision 4.61 with 400X magnification. The inflammatory process was evaluated by the pro-inflammatory cytokines (IL 1 and TNF α). Cytokine concentrations were measured in plasma collected from rats by multiplex immunoassay, using the MILLIPLEX MAP RatCytokine / Chemokine magnetic panel according to the manufacturer’s instructions (Millipore). The species’ cross-reactivity was evaluated by the manufacturer (http://www.abacusals.com/media/Milliplex_2014.pdf, Millipore system, MAGPIX® and MILLIPLEX® Analyst 5.1 software).

### 2.6 Statistics

The data were represented as the mean (X) ± standard deviation from the mean (SD). Differences of p <0.05 were considered significant. The results were expressed as χ ± SD. The significance of the differences between the means was analyzed by the Student’s t test. A value of p <0.05 was considered significant. GraphPad Prism 6.0 (GraphPad Software, La Jolla, California, United States) was used to calculate the p values of the difference.

## 3 RESULTS AND DISCUSSION

### 3.1 Hematoxylin and Eosin (HE)

We found that the cardiac fibers were more widely spaced and had fewer nuclei. We also found a greater presence of fibroblasts in the UChB rats. We noticed a greater presence of red blood cells and a closer approximation between the cardiomyocytes nuclei in the UChBC.

**Fig.1.**
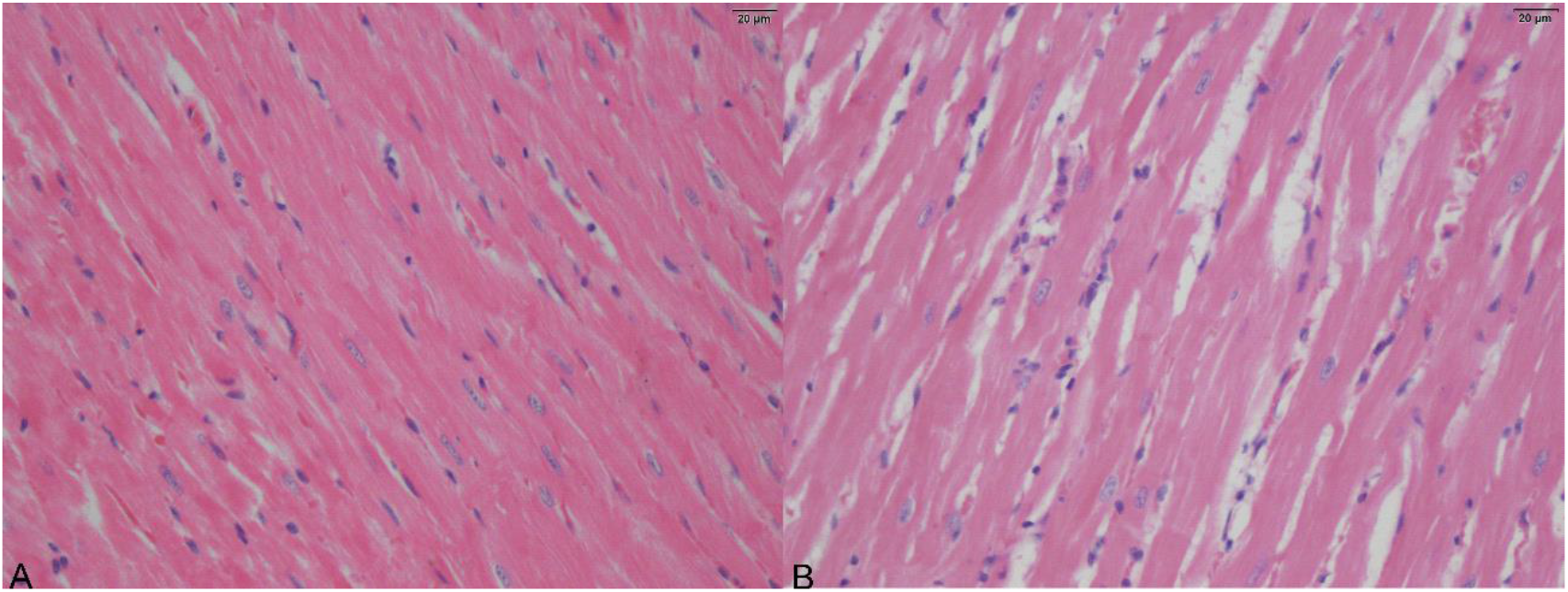
Sectional images of the left ventricle of UChBC (A) and UChB (B) rats. Both stained with HE.

**Fig. 2.**
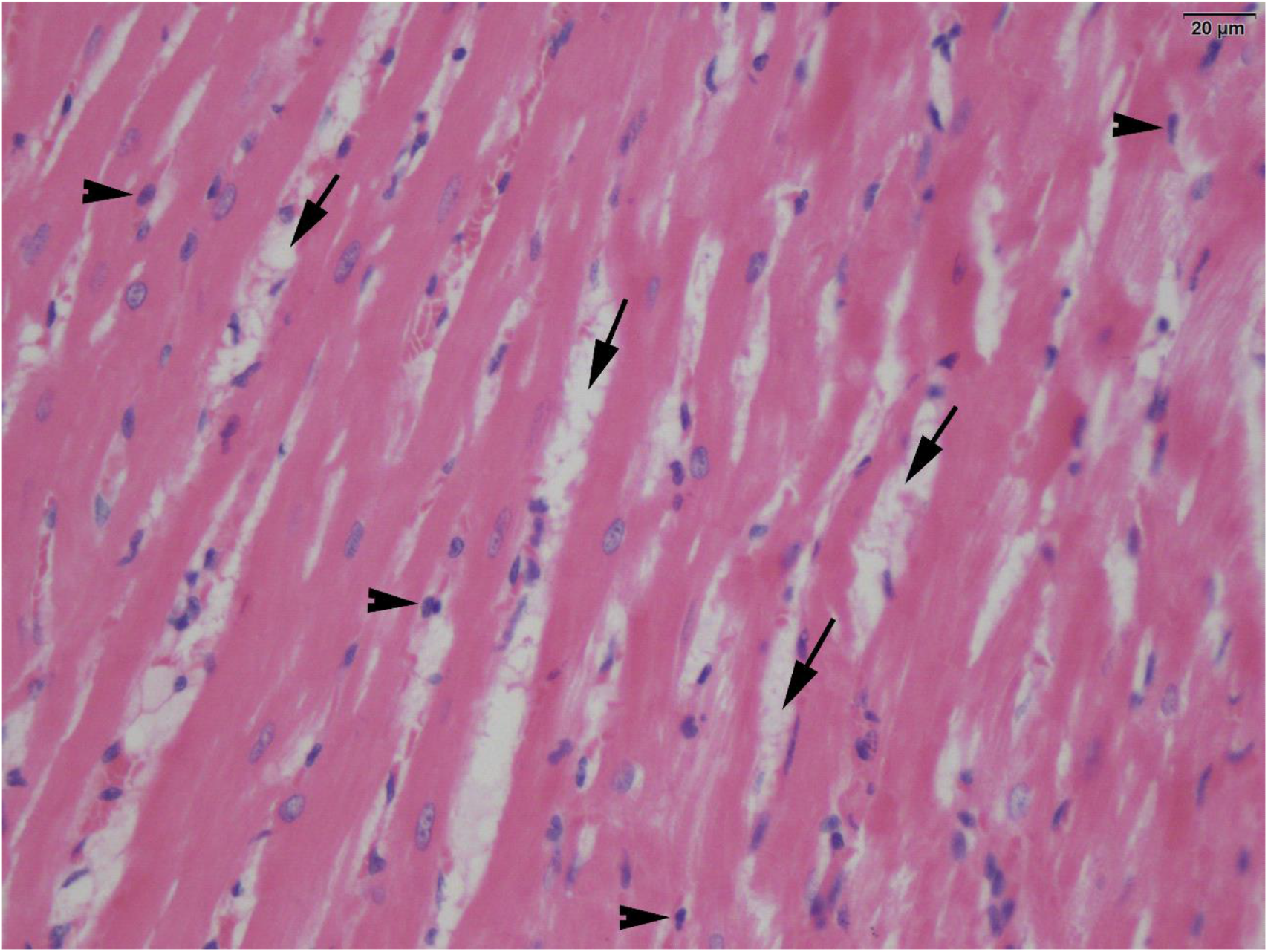
A sectional view of the left ventricle of the UChB. Arrows indicate the increased spaces between the fiber bundles (cardiomyocytes) and the arrowheads, the fibroblast nuclei. HE.

### 3.2 Masson’s trichrome (MT)

We found that the striations of the cardiac fibers are more evident in the UChBC, as the UChB presented myocytolysis. We noticed a greater presence of deposition of collagen fiber bundles in the UChB. In addition to changes linked to tissue remodeling, vascular changes also stood out, with the UChBs having less irrigation.

**Fig. 3.**
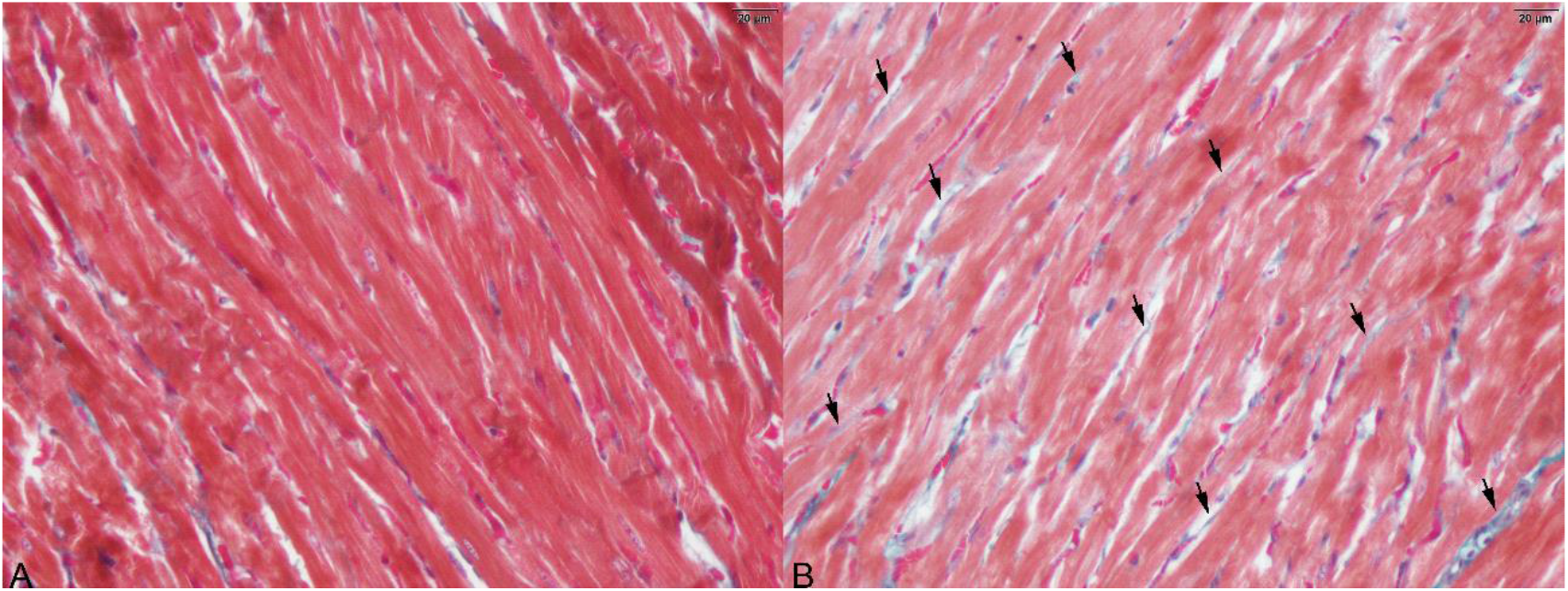
Sectional images of the left ventricle of UChBC (A) and UChB (B) rats. Arrows point at the bundles of collagen fibers. Both stained with MT.

**Fig. 4.**
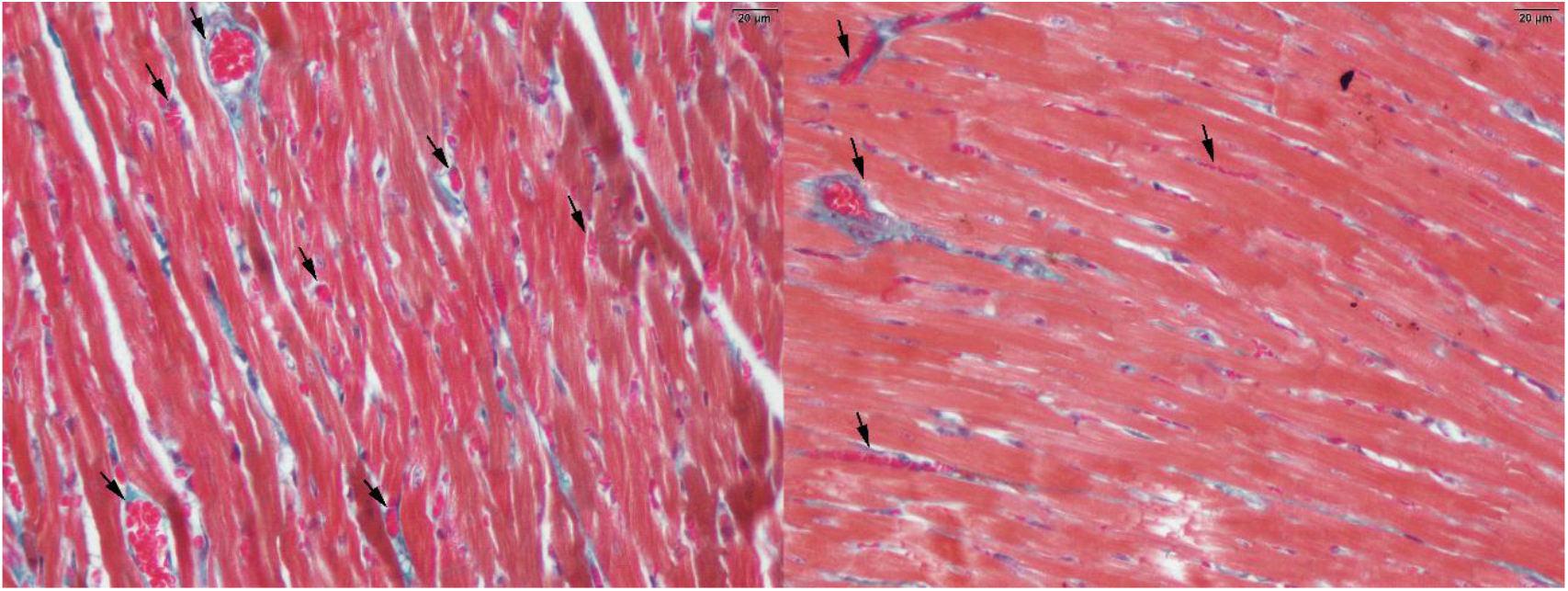
Sectional images of the left ventricle of UChBC (A) and UChB (B) rats. Arrows point to the vessels. Both stained with MT.

### 3.3 Ki67

We verified a greater proliferative activity in UChB in the immunohistochemical test for cell proliferation.

**Fig. 5.**
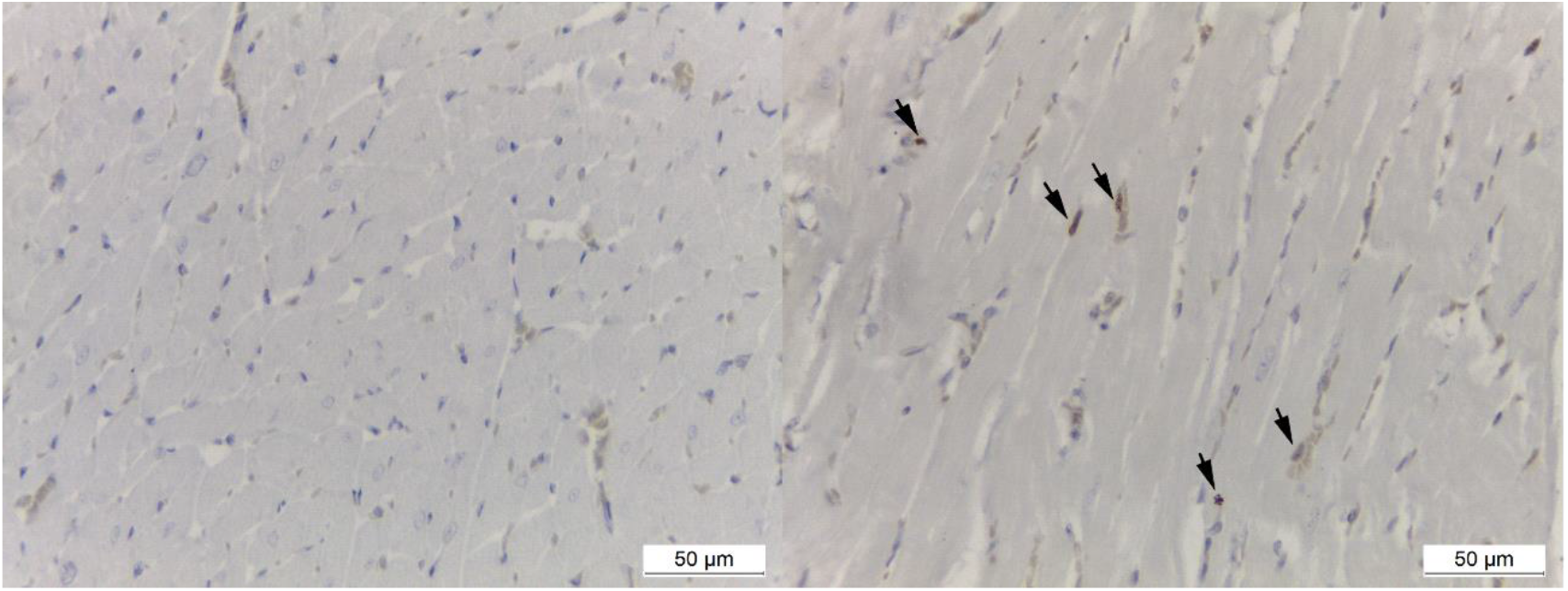
Sectional images of the left ventricle of UChBC (A) and UChB (B) rats. Arrows point to marked nuclei of proliferating cells.

### 3.4 Caspase 3

We verified a greater quantity of nuclei marked in the UChB in the immunohistochemical test referring to apoptosis.

**Fig. 6.**
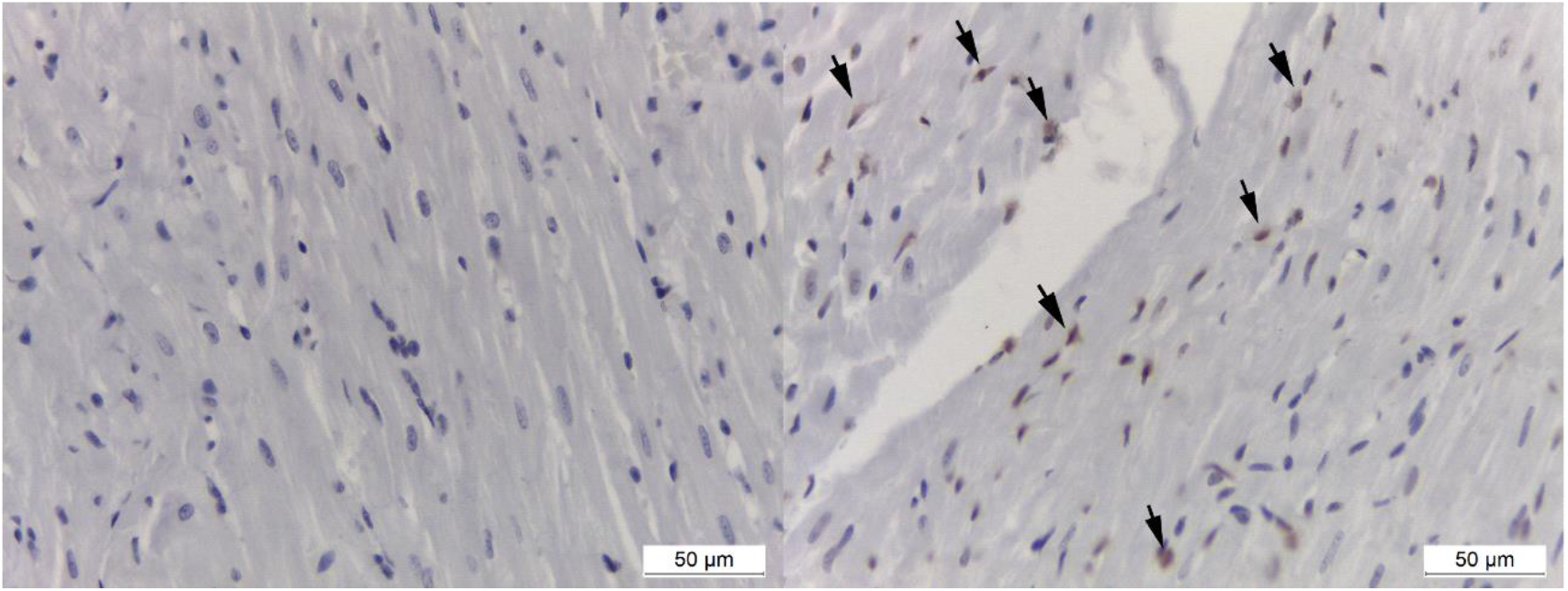
Sectional images of the left ventricle of UChBC (A) and UChB (B) rats. Arrows point to marked nuclei of cells undergoing apoptosis.

### 3.5 IL 1

The cytokine IL 1 showed no difference between groups.

**Fig. 7.**
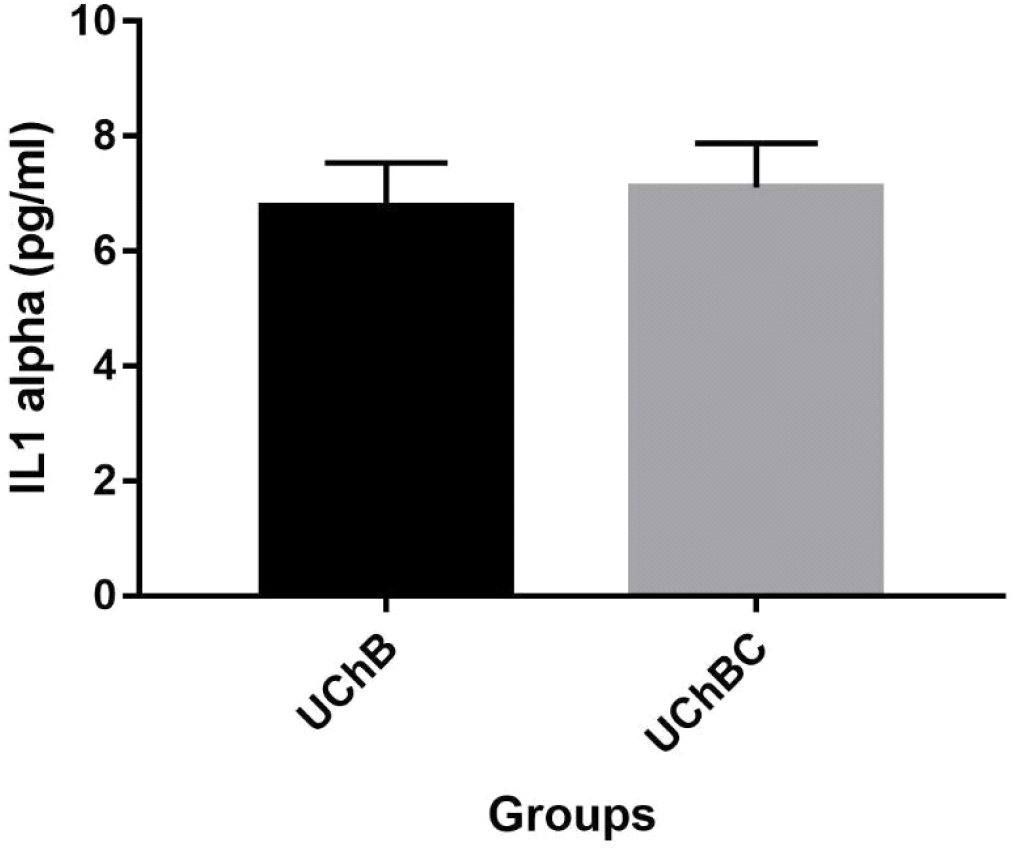
IL 1 alpha of the UChB and UChBC groups (data presented as mean ± SEM, n = 10 rats / group, p <0.05, unpaired T test)

### 3.6 TNF α

The TNF α cytokine showed no difference between groups.

**Fig. 8.**
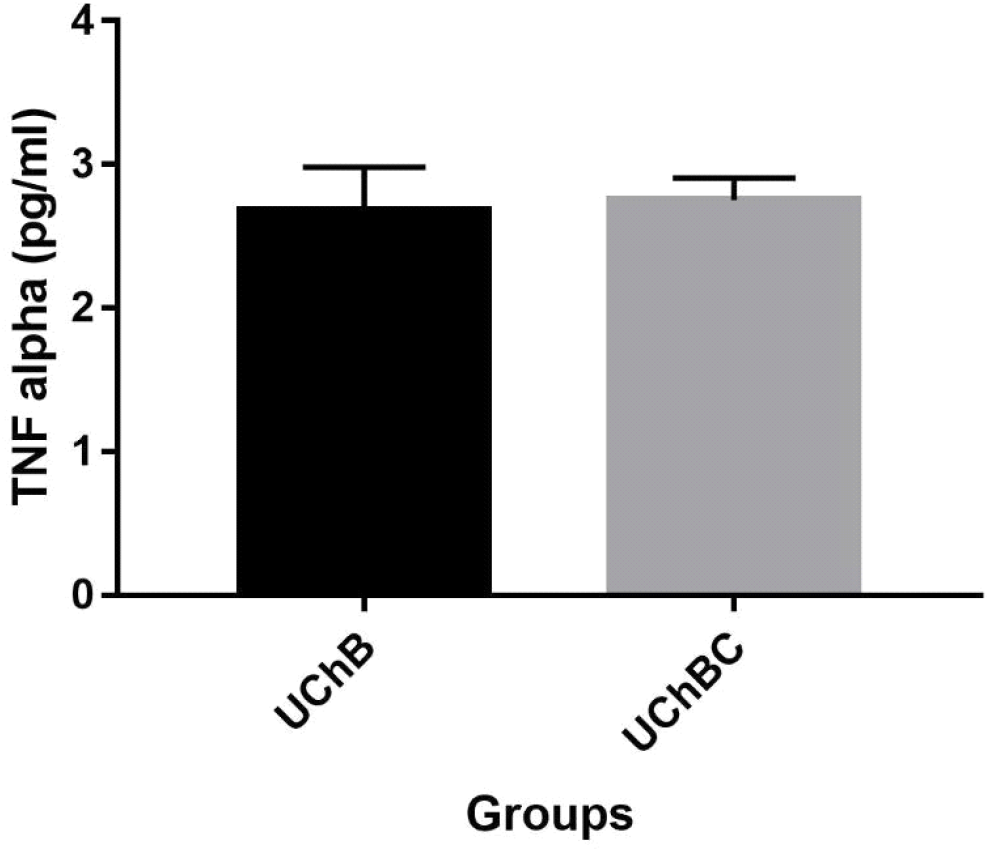
TNF alpha of the UChB and UChBC groups (data presented as mean ± SEM, n = 10 rats / group, p <0.05, unpaired T test)

The use of ethanol increases spacing between cardiac fibers, fewer nuclei, rise presence of fibroblasts, change the striations, up the deposition of bundles of collagen fibers and vascular changes. Ethanol is a biologically small and highly reactive molecule that easily diffuses into biological membranes and intracellular compartments, being able to hit all organelles. It interacts with membrane phospholipids, ion channels, and receptors, modifying their structure and function, changing intracellular transit, cellular energy and oxidative state [9,10]. In this injury environment, cardiomyocytes develop repair mechanisms such as hypertrophy and degrees of cell replacement, which can somehow mitigate the generation of injuries. Eventually, irreversible structural lesions develop with apoptosis.

The balance between mechanisms of cell proliferation and repair of lesions defines the degree and reversibility of cardiac damage [9]. The development of hypertrophy occurs due to the stress generated in the fibers and it occurs together with an increase in the montage of sarcomeres in response to the high demand for cardiac output. This effect is probably caused by increased circulating of growth factors and cytokines, along with neurohormonal activation [11,12]. The increase in the cellular volume of cardiomyocytes allows a smaller amount of fibers to occupy the place where there was a greater number of them before.

There is an increase in cardiac myostatin (Mstn) expression in alcoholic cardiomyopathy. Mstn inhibits mitogen-activated protein kinase (MAPK) [13,14] and interacts with metabolic proteins and enzymes [15]. MAPK is part of the growth factor signaling pathway which regulates cardiomyocyte activities and it is related to hypertrophy and anti-apoptotic and survival mechanisms [16]. Apoptosis of cardiomyocytes is the principal mechanism of cell loss induced by alcohol [17,18].

In addition to cellular and nuclear hypertrophy, the heart exposed to ethanol has myocytolysis and interstitial fibrosis. Myocytolysis is characterized by the dissolution and breakdown of myofibrils and cell vacuolization [9]. This can create changes in the interstice, affecting the metabolism and performance of cardiomyocytes and, ultimately, the function of the ventricles [19]. The observation of the increased spacing between the fiber is indicative of changes in the intercellular space. Possibly, the largest space is related to interstitial fibrosis, which increases the oxygen diffusion distance, reducing the partial arterial pressure of oxygen for cardiomyocytes. In addition, the electrical coupling of cardiomyocytes can be impaired by the accumulation of proteins and fibroblasts in the extracellular matrix [20].

Cardiomyocytes are mainly responsible for cardiac contractility, however they represent only 30% of healthy heart cells in rats and 28% in humans. Cardiac fibroblasts represent 64% of cardiac cells in rats and 72% in humans [21,22]. Thus, the increase in the number of fibroblasts is worrying, as they act in the processes of hypertrophy and fibrosis. Fibroblast growth factor 2 (Fgf-2) is the main paracrine promoter in the induction of cardiac hypertrophy through the MAPK activation pathway [23,24]. Fibroblast is the main source of Fgf-2 in the heart [25]. Fgf-2 has isoforms, one of high molecular weight and one of low, both of which are produced by the same mRNA [26,27]. High molecular weight Fgf-2 induces cardiac hypertrophy in vivo [28] (JIANG et al., 2007) and cardiac fibroblasts produce only this high molecular weight protein [25] through stimulation of angiotensin II and secrete by caspase-1 [29].

There is no inflammatory process characterized by the pro-inflammatory cytokines IL 1 and TNF, possibly due to the adaptive process of chronic use of ethanol in the variety. However, other injury mechanisms may be present, such as an imbalance in cellular metabolism.

## 4 CONCLUSION

Ethanol alters the myocardium structure and metabolism of the voluntary ethanol drinking rats’ left ventricle, impairing cardiac function. Our findings arouse interest in deepening the mechanisms involved.

## GLOSSARY

AC: alcoholic cardiomyopathy
CNS: central nervous system
UChB rat: University of Chile, consumption greater than 2 g of ethanol / Kg body weight/day, rats
HE: Hematoxylin & Eosin
MT: Masson’s Trichrome
Mstn: myostatin
MAPK: mitogen-activated protein kinase
Fgf-2: Fibroblast growth factor 2

Declarations of interest: none

## 5 ACKNOWLEDGMENT

Grant 2018/12354-5, 2018/26294-4 São Paulo Research Foundation (FAPESP)

